# Ventral posterolateral and ventral posteromedial thalamus neurons have distinct synaptic and cellular physiology

**DOI:** 10.1101/2022.12.30.521121

**Authors:** Carleigh Studtmann, Marek Ladislav, Mona Safari, Yang Chen, Mackenzie A. Topolski, Rabeya Khondaker, Eni Tomović, Aleš Balík, Sharon A. Swanger

**Affiliations:** Fralin Biomedical Research Institute at VTC, Center for Neurobiology Research, Roanoke, VA, USA; Graduate Program in Translational Biology, Medicine, and Health, Virginia Polytechnic and State University, Blacksburg, VA, USA; Institute of Physiology, Czech Academy of Sciences, Prague, Czech Republic; Department of Biomedical Sciences and Pathobiology, Virginia-Maryland College of Veterinary Medicine, Virginia Polytechnic and State University, Blacksburg, VA, USA; Department of Internal Medicine, VTC School of Medicine, Roanoke, VA, USA

## Abstract

Somatosensory information is propagated from the periphery to the cerebral cortex by two parallel pathways through the ventral posterolateral (VPL) and ventral posteromedial (VPM) thalamus. VPL and VPM neurons receive somatosensory signals from the body and head, respectively. VPL and VPM neurons also receive cell-type-specific GABAergic input from the reticular nucleus of the thalamus (nRT). Although VPL and VPM neurons have distinct connectivity and physiological roles, differences in the functional properties of VPL and VPM neurons remain unclear as they are often studied as one ventrobasal (VB) thalamus neuron population. Here, we directly compared synaptic and intrinsic properties of VPL and VPM neurons in C57Bl/6J mice of both sexes aged P25-P32. Recordings of spontaneous synaptic transmission suggested that VPL neurons receive excitatory synaptic input with higher frequency and strength than VPM neurons, while VPL neurons exhibited weaker inhibitory synapse strength than VPM neurons. Furthermore, VPL neurons showed enhanced depolarization-induced spike firing and greater spike frequency adaptation than VPM neurons. VPL and VPM neurons fired similar numbers of spikes during hyperpolarization rebound bursts, but VPM neurons exhibited shorter burst latency compared to VPL neurons, which correlated with increased sag potential during hyperpolarization. This work indicates that VPL and VPM thalamocortical neurons are functionally distinct populations. The observed functional differences could have important implications for their specific physiological and pathophysiological roles within the somatosensory thalamocortical network.

## Introduction

The corticothalamic (CT) network comprises a series of reciprocal loops between the cerebral cortex and thalamus that play essential roles in sensory, motor, emotional, and cognitive processing. The somatosensory CT circuit propagates somatic information from the periphery to the cortex through the ventrobasal (VB) thalamus and mediates oscillatory activity that regulates attention and sleep-wake cycles (Beenhakker & Huguenard, 2009; Fanselow & Nicolelis, 1999; Fernandez & Luthi, 2020; Wolff & Vann, 2019). Notably, somatic information from the body and the head are processed via two parallel pathways involving the ventral posterolateral (VPL) and ventral posteromedial (VPM) regions of the VB thalamus. These two somatosensory pathways make distinct contributions to sensory processing and oscillatory activity; however, the synaptic and cellular mechanisms underlying pathway-specific physiology and pathophysiology are not fully understood.

VPL neurons receive ascending somatosensory information from the body via the medial lemniscus and spinothalamic tract, whereas somatosensory information from the head is propagated to VPM neurons via the trigeminothalamic tract (O’Reilly et al., 2021). Thalamocortical neurons in the VPL and VPM relay somatosensory information to the primary somatosensory cortex (S1), and layer 6 CT neurons return glutamatergic feedback to the VPL and VPM (Aziz & Ahmad, 2006; Brecht & Sakmann, 2002; Lenz, 1992). CT neurons as well as VPL and VPM thalamocortical neurons send collaterals to the reticular nucleus of the thalamus (nRT), a sheet of GABAergic neurons that provide the primary inhibitory input to VPL and VPM neurons (**Figure 1A**). Interestingly, VPL neurons receive input from both somatostatin (SOM)- and parvalbumin (PV)-expressing nRT neurons, while VPM neurons receive input from only PV-expressing nRT neurons (Clemente Perez et al., 2017). Both VPL and VPM neurons exhibit tonic and burst firing modes important for processing somatosensory information and generating intra-thalamic oscillations through reciprocal connections with the nRT (Llinas & Steriade, 2006; Sherman, 2001; Steriade & Llinas, 1988). However, the nRT-VPM loop is more rhythmogenic than the nRT-VPL loop; this difference may be due to the more robust connections between VPM neurons and PV-expressing nRT neurons or distinct intrinsic properties of VPL and VPM neurons. While numerous studies have investigated the physiological properties of the VB thalamus as a whole, it remains unclear if VPL and VPM neurons have differences in synaptic and cellular function that contribute to their distinct physiological roles (Abbas et al., 2006; Astori et al., 2011; Cheong et al., 2011; Ha et al., 2016; Huguenard & Prince, 1994; Jacobsen et al., 2001; Talley et al., 1999; Warren et al., 1994; Zobeiri et al., 2019).

**Figure 1.**
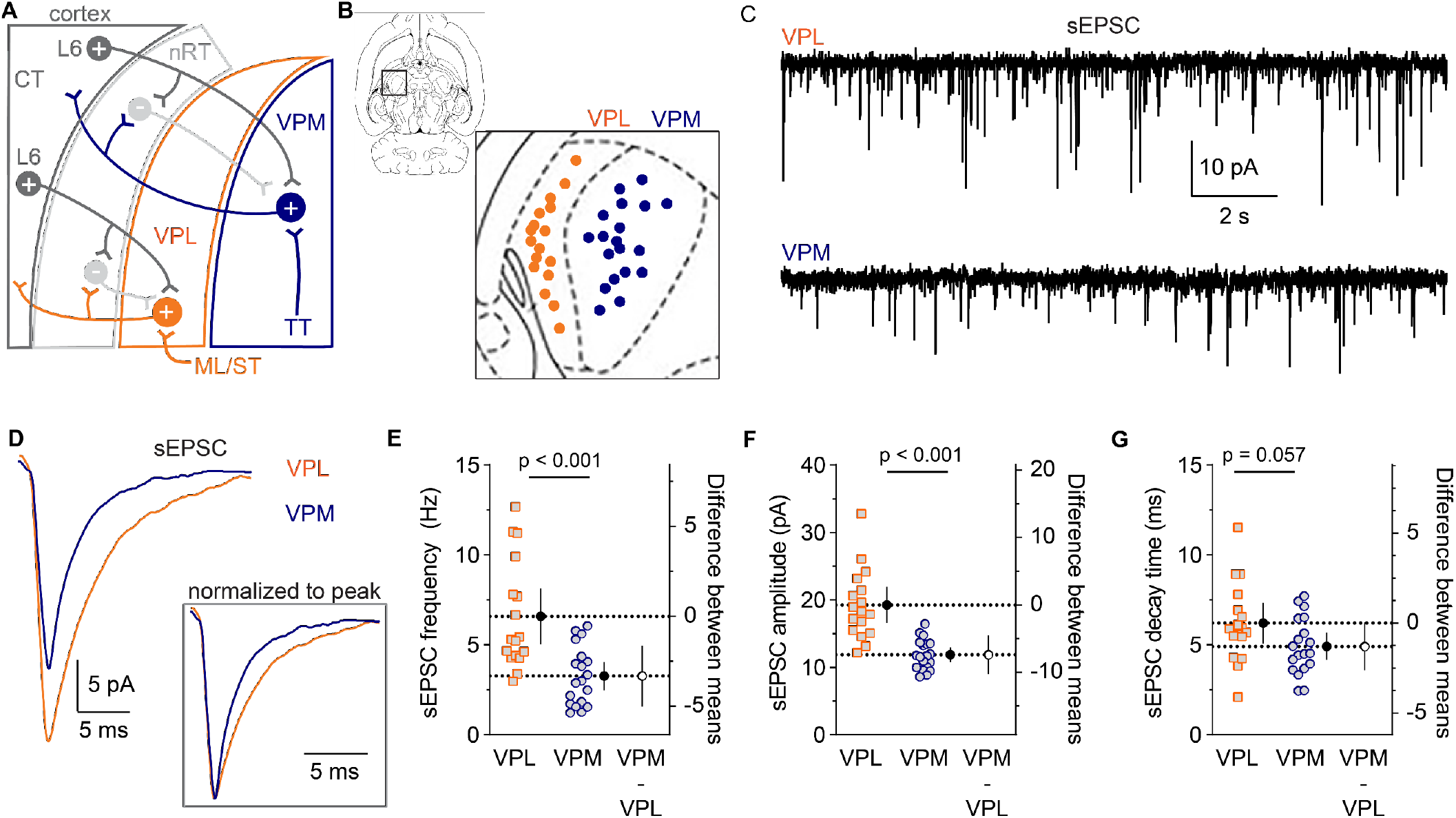
sEPSC frequency and amplitude were reduced in VPM neurons compared to VPL neurons. **A.** A somatosensory CT circuit diagram shows glutamatergic (+) and GABAergic (-) neuron connectivity as well as the ascending somatosensory inputs via the trigeminothalamic tract (TT), medial lemniscus (ML), and spinothalamic tract (ST). **B.** An image from a horizontal rat brain atlas (DV -3.1 mm relative to bregma) was used to map the cell location for each sEPSC recording. VPL: *n* = 17 cells, from 11 mice. VPM: *n* = 19 cells, 12 mice. Current traces from representative VPL and VPM neurons show (**C**) sEPSC voltage-clamp recordings and (**D**) ensemble averages of all sEPSCs. Inset in panel D: sEPSCs were normalized to the peak response. Welch’s t tests were used to compare sEPSC (**E**) frequency (*t* = 4.043, *df* = 23.87), (**F**) amplitude (*t* = 5.443, *df* = 21.24), and (**G**) decay time (*t* = 1.985, *df* = 28.60), which were plotted on the left y-axis as individual data points for each cell alongside the group means (black circle, dashed line) with 95% CI. The difference between means (VPM – VPL) was plotted (open circle) with 95% CI on the right y-axis. P values were reported above each graph.

VB thalamus dysfunction, including pathological oscillatory activity, has been implicated in many diseases (David et al., 2018; Hall & Lifshiftz, 2010; Hazra et al., 2016; Jeanne T. Paz, 2013; Paz et al., 2011; Princivalle et al., 2003; Stefanie Ritter-Makinson, 2019; Ulrike B.S. Hedrich, 2014). However, the distinct connectivity and oscillatory activity of VPL and VPM neurons suggest the two neuron populations may differentially contribute to circuit dysfunction in disease. Indeed, VPM neuron activity has been proposed to make larger contributions to pathological oscillations during seizures, and the tonic and burst firing properties of VPL and VPM neurons were differentially affected in a mouse model of Dravet syndrome, an infantile epileptic encephalopathy (Clemente-Perez et al., 2017; Studtmann et al., 2022). In a spinal cord injury mouse model, VPL neurons exhibited a compensatory upregulation in the voltage-gated sodium channel Na_V_1.3, whereas VPM neurons did not (Peng Zhao, 2006). These disease states could impact VPL and VPM neuron firing or ion channel expression differently due to their distinct connectivity or unique intrinsic regulatory mechanisms. From a therapeutic perspective, if altered function in specific cell populations underlie circuit dysfunction, then effective treatments may require targeting those particular cells, not all cells within the circuit.

Intrinsic and synaptic properties of VPL and VPM neurons have been studied individually, but there has been no direct systematic comparison (Chiaia et al., 1991; Landisman & Connors, 2007). Because these two populations receive unique inputs and respond differentially in some disease states, we hypothesized that VPL and VPM neurons have unique synaptic and cellular physiology that contribute to their distinct roles in somatosensory CT circuit function. A thorough understanding of the synaptic and cellular properties of VPL and VPM neurons is essential to elucidate how they may be uniquely processing information and contributing to circuit function or dysfunction. Here, we report substantial differences in VPL and VPM neuron excitatory and inhibitory synaptic input as well as tonic and burst firing properties. These findings broaden our understanding of nucleus-specific functional properties within the thalamus and provide evidence that the VPL and VPM constitute physiologically distinct neuron populations.

## Materials and Methods

### Preparation of acute mouse brain slices

Mouse studies were performed according to protocols approved by the Institutional Animal Care and Use Committee at Virginia Polytechnic Institute and State University and in accordance with the National Institutes of Health guidelines. C57Bl6/J mice of both sexes aged P25-P32 were used for all experiments. Mice were housed in a 12-hour light/dark cycle with *ad libitum* access to food and water. Mice were deeply anesthetized with an overdose of inhaled isoflurane and transcardially perfused with ice-cold sucrose-based artificial cerebrospinal fluid (aCSF) containing (in mM) 230 sucrose, 24 NaHCO_3_, 10 glucose, 3 KCl, 10 MgSO_4_, 1.25 NaH_2_PO_4_, and 0.5 CaCl_2_ saturated with 95% O_2_ / 5% CO_2_. The brain was removed and glued to a vibratome stage (Leica VT1200S), and horizontal 300 μm slices were cut in an ice-cold sucrose aCSF bath. Slices were incubated in a NaCl-based aCSF containing (in mM) 130 NaCl, 24 NaHCO_3_, 10 glucose, 3 KCl, 4 MgSO_4_, 1.25 NaH_2_PO_4_, and 1 CaCl_2_ saturated with 95% O_2_ / 5% CO_2_ at 32°C for 30 min. The slices were then equilibrated to room temperature (RT) for 30 minutes and maintained at RT until used for recordings up to 10 hr later.

### Electrophysiology

One cell was recorded per brain slice, and no more than two cells in any dataset are from the same mouse. Recordings were made using a Multiclamp 700B amplifier (Molecular Devices), sampled at 20 kHz (Digidata 1550B, Molecular Devices), and low-pass filtered at 10 kHz using Axon pClamp 11 software (Molecular Devices). The extracellular recording solution contained (in mM) 130 NaCl, 24 NaHCO_3_, 10 glucose, 3 KCl, 1 MgSO_4_, 1.25 NaH_2_PO_4_, and 2 CaCl_2_ saturated with 95% O_2_ / 5% CO_2_, and was maintained at 32°C for all recordings. For whole-cell voltage-clamp recordings, borosilicate glass recording electrodes (4-5 MΩ) were filled with (in mM) 120 CsMeSO_3_, 15 CsCl, 8 NaCl, 10 tetraethylammonium chloride, 10 HEPES, 1 EGTA, 3 Mg-ATP, 0.3 Na-GTP, 1.5 MgCl_2_, 1 QX-314, and 0.1% biocytin, pH 7.3. After a 10-minute equilibration period, spontaneous excitatory postsynaptic currents (sEPSCs) and spontaneous inhibitory postsynaptic currents (sIPSCs) were recorded in two-minute epochs at a holding potential of -70 mV and 0 mV, respectively. Holding commands were adjusted for a 10 mV liquid junction potential during the recordings. Series resistance and cell capacitance were monitored throughout the experiment, but were not compensated, and cells were excluded if either parameter changed > 20%.

For whole-cell current-clamp recordings, borosilicate glass recording electrodes (4-5 MΩ) were filled with (in mM) 130 K-gluconate, 4 KCl, 2 NaCl, 10 HEPES, 0.2 EGTA, 4 ATP-Mg, 0.3 GTP-Tris, 14 phosphocreatine-K, and 0.1% biocytin, pH 7.3. Pipette capacitance neutralization and bridge balance were enabled during current-clamp recordings for capacitance and series resistance compensation. Membrane potential values were corrected for the liquid junction potential after the recording (15 mV). The resting membrane potential (RMP) was determined 2 min after breakthrough. To measure intrinsic membrane properties, voltage responses were elicited by 200 ms hyperpolarizing current injections between 20 – 100 pA (20 pA steps). Synaptic blockers APV (100 μM), NBQX (10 μM), and gabazine (10 μM) were washed into extracellular solution and applied for at least five minutes prior to current injection experiments. Depolarization-induced spike firing was elicited by 500 ms depolarizing current injections between 20 – 400 pA (20 pA steps). Hyperpolarization-induced rebound bursting was elicited by 500 ms hyperpolarizing current injections between 50 – 400 pA (50 pA steps). All current-clamp experiments were conducted from a resting membrane potential (RMP) of -75 ± 2 mV, which was maintained with a bias current (0 – 30 pA). Three trials were completed for each current injection experiment.

### Confirmation of cell location

The cell location in the VPL or VPM was confirmed after electrophysiology recordings by biocytin labeling as previously described (Studtmann et al., 2022) or by images of the electrode location during the recording, and then mapped onto a rodent brain atlas image (Gaidica, 2022). For biocytin labeling, brain slices were fixed with 4% paraformaldehyde in 1X PBS, pH 7.4, overnight at 4°C, washed in 1X PBS, and then stored at -20°C in cryoprotectant solution containing 0.87 M sucrose, 5.37 M ethylene glycol, and 10 g/L polyvinyl-pyrrolidone-40 in 0.1 M phosphate buffer, pH 7.4. Slices were blocked with 10% NDS in 1X PBS with 0.25% Triton X-100, and then incubated with 1.0 μg/ml DyLight 594-conjugated streptavidin (Jackson Immunoresearch) in blocking solution overnight at RT. Slices were mounted on a glass slide with a coverslip and DABCO mounting media. 10X images were acquired on an Olympus IX83 microscope with a Hamamatsu Orca Flash 4.0 camera, X-Cite Xylis LED, and TRITC filter sets using CellSens software.

### Electrophysiology data analysis

Recordings were assigned numerical identifiers and analysis was performed blind to cell location. For sIPSC and sEPSC analysis, recordings of 4 – 6 min were analyzed to determine interevent interval, amplitude, decay times, and burst properties using MiniAnalysis software (Synaptosoft). Data were digitally filtered at 1 kHz. Automated detection identified events with amplitude equal to or greater than 5 x RMS noise level, which was 8 – 12 pA, and then automated detection accuracy was assessed and any additional events ≥ 6 pA were detected manually. Amplitude and frequency values for each sEPSC or sIPSC recording were averaged in 30 second bins, and the reported values for each cell are the average across bins. The sEPSC and sIPSC decay times for each cell were determined from ensemble averages of all sEPSCs or sIPSCs recorded from each cell. The 10-90% peak to baseline decay times of the ensemble averages were fitted using the following double-exponential function:

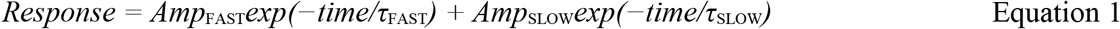

where τ_FAST_ is the fast deactivation time constant, τ_SLOW_ is the slow deactivation time constant, *Amp*_FAST_ is the current amplitude of the fast deactivation component, and *Amp*_SLOW_ is the current amplitude of the slow deactivation component. The weighted decay time constant (τW) was calculated by:

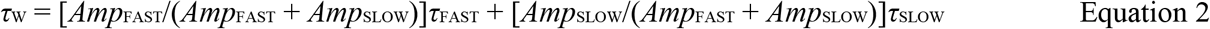

Current-clamp recordings were analyzed in Clampfit 11 (Molecular Devices). Input resistance (R_in_) was determined from the amplitude of voltage responses to 200 ms hyperpolarizing current injections, the time constant (τm) was determined by a mono-exponential fit of the voltage response, and cell capacitance was calculated by C_m_ = τ_m_/R_in_. The Clampfit 11 threshold detection module was used to quantify the number of spikes, spike frequency, and latency to the first spike in response to 500 ms depolarizing current injections or upon removal of 500 ms hyperpolarizing current injections. Rheobase was defined as the smallest depolarizing current injection that elicited an action potential. Values were averaged across three runs for all current injection experiments.

The shape of single action potentials or low frequency trains of action potentials were analyzed using the Action Potential Search module in ClampFit 11. The baseline was manually set, and action potential amplitude, half width, rise and decay time, after-hyperpolarization amplitude and duration, and threshold were measured. Spike frequency adaptation was evaluated by measuring spike frequency across depolarizing current injections. Some VPL and VPM neurons had a prolonged inter-event interval between the first two spikes following depolarization; therefore, the frequency of the first three spikes following current injection (*f*_start_) was measured in VPL and VPM neurons and compared to the frequency of the last two spikes (*f*_end_). The ratio of these frequencies (*f*_start_ / *f*_end_) is reported as the adaptation ratio. The frequencies from two or three runs in the same cell were averaged.

### Statistical analysis

*A priori* power analyses were performed in GPower 3.1 to estimate required samples sizes given appropriate statistical tests with α = 0.05, power (1 – β) = 0.8, and a moderate effect size or effect size based on pilot data. Statistical analyses were performed using GraphPad Prism 9.4.1. All datasets were determined to be normally distributed by the Shapiro-Wilk test (*p* > 0.05). Group data are plotted in estimation plots including the individual data points for each cell and the group mean with 95% confidence interval (CI) on the left y-axis as well as the difference between means (*M*_VPM_ - *M*_VPL_) with 95% CI on the right y-axis. Data are reported in the text as the mean with standard deviation (*SD)* as well as the difference between means with 95% CI. Negative values reflect a decrease in *M*_VPM_ relative to *M*_VPL_, and positive values reflect an increase in *M*_VPM_ relative to *M*_VPL_. Equal variances between groups were not assumed as previous data showed unequal variances between VPL and VPM neuron populations for several metrics (Studtmann et al., 2022). As such, pairwise comparisons of group means were performed with Welch’s t tests. Mixed-effects analysis was used for repeated measures current-clamp recording data where measurements were analyzed across multiple current injections or spike trains within each sample. The main effect of group (VPL vs. VPM) was the primary statistical outcome, and thus the overall mean across current injections or spike trains was reported with *SD* in the text and graphed in estimation plots with 95% CI. The interaction effect was also reported in the figure legends. We did not perform pairwise comparisons for specific current injection levels or spike pairs. Correlations were analyzed by computing Pearson correlation coefficients, and the data are reported in the text as *r* with 95% CI and *R*^2^. The data were plotted and fit with a straight line using the least squares regression method. All statistical inferences with exact p values are reported in the text or tables as well as the figures. The specific statistical tests and sample sizes are reported in the figure legends or table captions.

## Results

### VPL neurons have enhanced excitatory synaptic input compared to VPM neurons

VPL and VPM neurons receive somatosensory information via anatomically distinct glutamatergic inputs, which could exhibit differences in synaptic transmission (**Figure 1A**). We compared excitatory synaptic input to VPL and VPM neurons by recording sEPSCs in acute brain slices from C57Bl/6J mice aged P25-P32 (**Figure 1B – D)**. The mean sEPSC frequency in VPM neurons (3.3 Hz, *SD* 1.6) was reduced by 3.3 Hz [-5.0, -1.6] compared to VPL neurons (6.6 Hz, *SD* 3.0; *p* < 0.001; **Figure 1E**). The mean sEPSC amplitude in VPM neurons (11.9 pA, *SD* 2.2) was reduced by 7.4 pA [-10.2, -4.6] compared to VPL neurons (19.3 pA, *SD* 5.2,*p* < 0.001; **Figure 1F**). The mean sEPSC decay time in VPM neurons (4.9 ms, *SD* 1.6) was 1.3 ms [-2.6, 0.0] less than VPL neurons (6.2 ms, *SD* 2.2, *p* = 0.057; **Figure 1G**). These findings suggest that VPM neurons have reduced frequency and strength of glutamatergic synaptic input compared to VPL neurons.

### VPM neurons have higher amplitude inhibitory synaptic input than VPL neurons

Recent work indicates that PV-positive nRT neurons project to both VPL and VPM neurons, whereas SOM-positive neurons project to VPL neurons, but not VPM neurons (**Figure 2A**)(Clemente-Perez et al., 2017). To test if this distinct connectivity manifests in differences in inhibitory synaptic input, we measured the frequency, strength, and decay kinetics of spontaneous inhibitory postsynaptic currents (sIPSCs) in VPL and VPM neurons. The mean sIPSC frequency was 4.9 Hz (*SD* 2.3) in VPM neurons and 6.9 Hz (*SD* 3.1) in VPL neurons, whereas sIPSC amplitude was 29.5 pA (*SD* 3.6) in VPM neurons and 25.9 pA (*SD* 3.7) in VPL neurons (**Figure 2B-D**). These data indicate that the sIPSC frequency was 1.9 Hz [-3.9, 0.0] lower in VPM neurons compared to VPL neurons (*p* = 0.058), and sIPSC amplitude was increased by 3.6 pA [1.3, 5.9] in VPM neurons compared to VPL neurons (*p* = 0.003; **Figure 2E,F)**. VPM neuron mean sIPSC decay time (4.4 ms, *SD* 0.7) was 0.3 ms [-0.8, 0.1] less than VPL neurons (4.7 ms, *SD* 0.7, *p* = 0.127; **Figure 2G**). These data suggest that VPM neurons received higher amplitude sIPSCs than VPL neurons, but sIPSC frequency and decay time were similar between the two neuron populations.

**Figure 2.**
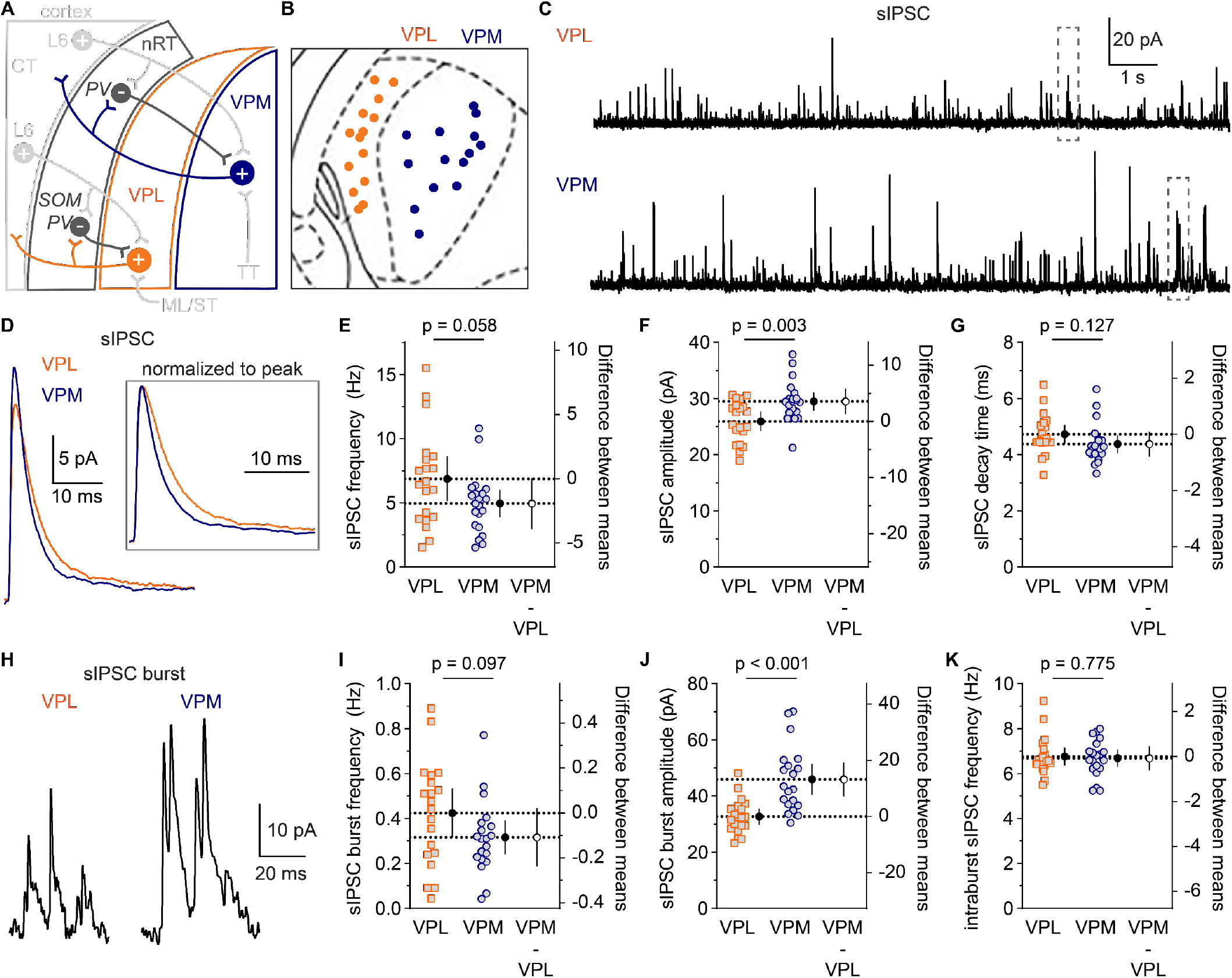
sIPSC amplitude was higher in VPM neurons than VPL neurons. **A.** The CT circuit diagram illustrates PV- and SOM-positive GABAergic nRT neuron connectivity (dark gray) with VPL and VPM neurons. **B.** A map of the cell location for each sIPSC recording. VPL: *n* = 20, 14 mice. VPM: *n* = 21 cells, 15 mice. Current traces from representative VPL and VPM neurons show (**C**) sIPSC voltage-clamp recordings and (**D**) ensemble averages of all sIPSCs. Inset in panel D: ensemble average sIPSCs were normalized to the peak response. Welch’s t test were used to compare sIPSC (**E**) frequency (*t* = 1.970, *df* = 31.76), (**F**) amplitude (*t* = 3.155, *df* = 38.85), and (**G**) decay time (*t* = 1.559, *df* = 38.81), which were plotted on the left y-axis as individual data points for all cells alongside the group mean (black circle, dashed line) with 95% CI. The difference between means (VPM – VPL) was plotted (open circle) with 95% CI on the right y-axis. **H**. Representative traces show large amplitude bursts of sIPSCs in VPL and VPM neurons that were expanded from boxed regions in panel C. Welch’s t tests were used to compare sIPSC (**I**) burst frequency (*t* = 1.709, *df* = 33.99), (**J**) burst amplitude (*t* = 4.475, *df* = 28.77), and (**K**) intraburst frequency (*t* = 0.288, *df* = 37.91), which were plotted as described for panels E-G. VPL: *n* = 20 cells, 14 mice. VPM: *n* = 20 cells, 15 mice. P values were reported above each plot.

PV-positive nRT neurons exhibit stronger burst firing properties than SOM-positive nRT neurons (Clemente-Perez et al., 2017), which suggests that bursts of inhibitory input to VPL and VPM neurons may differ. On average, sIPSC burst frequency in VPM neurons (0.32 Hz, *SD* 0.16) was 0.1 Hz [-0.24, 0.02] lower than VPL neurons (0.42 Hz, *SD* 0.23, *p* = 0.097; **Figure 2H,I**). The mean sIPSC burst amplitude was 13.2 pA [7.2, 19.3] higher in VPM neurons (45.9 pA, *SD* 12) than VPL neurons (32.7 pA, *SD* 6.2,*p* < 0.001; **Figure 2J**). The mean intraburst sIPSC frequency was 6.7 Hz (*SD* 0.8) for VPM neurons and 6.8 Hz (*SD* 0.9) for VPL neurons, which is a mean difference of -0.1 Hz [-0.6, 0.5], *p* = 0.775 (**Figure 2K**). These data suggest that VPM neurons have significantly higher amplitude sIPSC bursts relative to VPL neurons, whereas sIPSC burst frequency was highly variable among neurons and showed no clear differences between groups and intraburst sIPSC frequency was similar between VPL and VPM neurons.

### Action potential afterhyperpolarization period differs between VPL and VPM neurons

To determine if cellular properties differed between VPL and VPM neurons, we compared intrinsic membrane properties and action potential (AP) shape. Intrinsic membrane properties were measured from the voltage response to small hyperpolarizing current injections (**Figure S1A,B**). The mean RMP, input resistance (Rin), cell capacitance (Cm), and membrane time constant (τ) were similar for VPL and VPM neurons (**Table 1; Figure S1C-F**) suggesting that their overall membrane properties were not different. AP shape parameters were measured from single APs or low frequency spike trains elicited at or near the rheobase current (**Figure 3A**). The effect sizes for AP threshold, rise slope, decay slope, half-width, and amplitude data all indicated that there were no substantial differences between VPL and VPM neurons (**Table 2; Figure 3B-E**). Interestingly, the afterhyperpolarization (AHP) amplitude and AHP duration were reduced in VPM neurons compared to VPL neurons (**Table 2; Figure 3C, F, G**). These data suggest that the rising and recovery phases of the AP are similar between VPL and VPM neurons, but distinct mechanisms may regulate the AHP period in the two neuron populations.

**Table 1.**
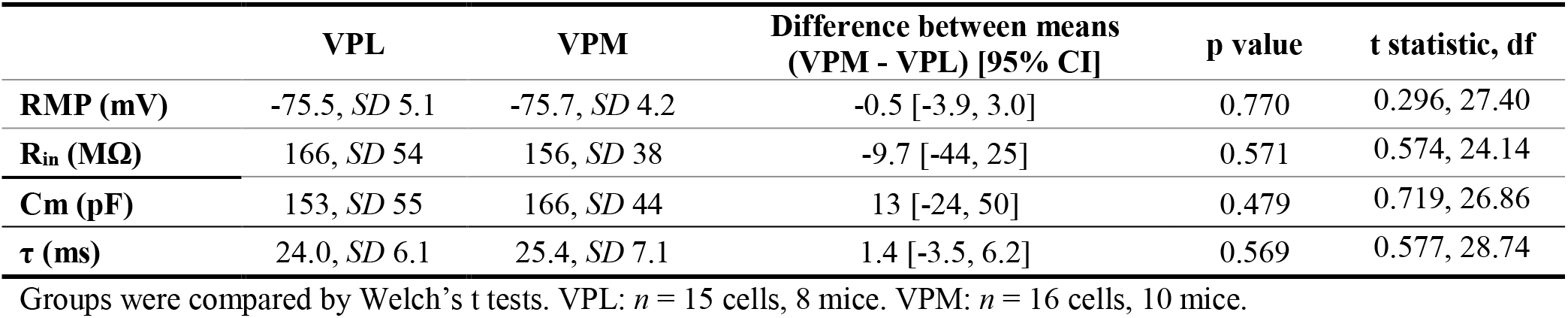
Intrinsic membrane properties of VPL and VPM neurons.

|  | VPL | VPM | Difference between means<br>(VPM - VPL) [95% CI] | p value | t statistic, df |
| --- | --- | --- | --- | --- | --- |
| **RMP (mV)** | -75.5, *SD* 5.1 | -75.7, *SD* 4.2 | -0.5 [-3.9, 3.0] | 0.770 | 0.296, 27.40 |
| **R<sub>in</sub> (MΩ)** | 166, *SD* 54 | 156, *SD* 38 | -9.7 [-44, 25] | 0.571 | 0.574, 24.14 |
| **Cm (pF)** | 153, *SD* 55 | 166, *SD* 44 | 13 [-24, 50] | 0.479 | 0.719, 26.86 |
| **τ (ms)** | 24.0, *SD* 6.1 | 25.4, *SD* 7.1 | 1.4 [-3.5, 6.2] | 0.569 | 0.577, 28.74 |

Groups were compared by Welch's t tests. VPL: *n* = 15 cells, 8 mice. VPM: *n* = 16 cells, 10 mice.

**Figure 3.**
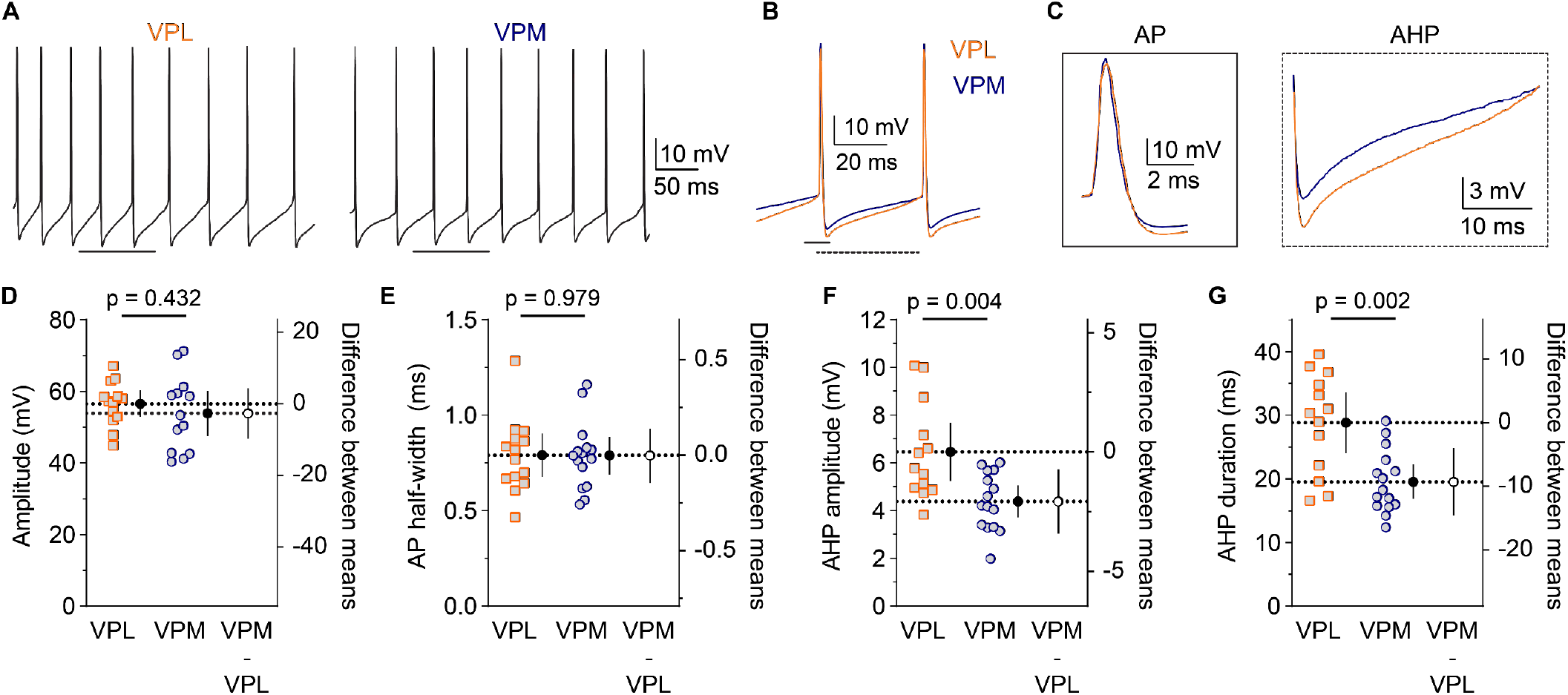
VPL neurons have an enhanced AHP period relative to VPM neurons. **A.** AP shape was determined from fitting single or low frequency AP trains elicited by depolarizing current injections in VPL and VPM neurons. **B**. The AP recording periods above the black lines in panel A were expanded and overlaid to illustrate shape differences. **C**. The left (solid black box) and right (dashed black box) panels were expanded from the region above the solid and dashed lines in panel B to show AP shape and the AHP period, respectively. (**D**) AP amplitude, (**E**) AP half-width, (**F**) AHP amplitude, and (**G**) AHP duration for all cells were plotted on the left y-axis as individual data points alongside the group mean (black circle, dashed line) with 95% CI. The difference between means (VPM – VPL) was plotted (open circle) with 95% CI on the right y-axis. Groups were compared with Welch’s t tests. P values are reported above each plot. See Table 2 for additional statistics.

**Table 2.** Action potential shape analysis of VPL and VPM neurons.

|  | VPL | VPM | Difference between means<br>(VPM - VPL) [95% CI] | p value | t statistic, df |
| --- | --- | --- | --- | --- | --- |
| **Threshold (mV)** | -46.2, *SD* 4.2 | -47.7, *SD* 3.7 | -1.5 [-4.7, 1.6] | 0.331 | 0.987, 24.92 |
| **Amplitude (mV)** | 56.6, *SD* 6.2 | 53.9, *SD* 10.6 | -2.7 [-9.8, 4.4] | 0.432 | 0.803, 19.40 |
| **Half-width (ms)** | 0.79, *SD* 0.19 | 0.79, *SD* 0.18 | 0.00 [-0.14, 0.14] | 0.979 | 0.027, 26.30 |
| **Rise slope (mV/ms)** | 138, *SD* 31 | 141, *SD* 51 | 3 [-33, 40] | 0.858 | 0.182, 18.15 |
| **Decay slope (mV/ms)** | -76, *SD* 11 | -71, *SD* 24 | 5 [-11, 21] | 0.527 | 0.647, 15.30 |
| **AHP amplitude (mV)** | 6.5, *SD* 2 | 4.4, *SD* 1.2 | -2.1 [-3.4, -0.7] | 0.004 | 3.232, 18.98 |
| **AHP duration (ms)** | 29, *SD* 8 | 20, *SD* 5 | -9 [-15, -4] | 0.002 | 3.684, 19.61 |

Groups were compared by Welch's t tests. VPL: *n* = 13 cells, 8 mice. VPM: *n* = 15 cells, 9 mice.

### Enhanced depolarization-induced firing in VPL compared to VPM neurons

We compared VPL and VPM neuron excitability by evaluating AP firing in response to a series of 500 ms depolarizing current injections (**Figure 4A**). The number of spikes fired at each current injection were quantified, and a mixed-effects model was used to analyze the overall mean spike number across all current injections (**Figure 4B**). VPM neurons fired an average of 18 spikes (*SD* 11), whereas VPL neurons fired an average of 28 spikes (*SD* 10), which is a mean difference of -10 spikes [-17, -2] (*p* = 0.012; **Figure 4B,C**). The mean rheobase current for VPM neurons (154 pA, *SD* 55) was 73 pA [35, 110] more than VPL neurons (81 pA, *SD* 52, *p* < 0.001; **Figure 4D**). The mean spike latency at the rheobase current was 98 ms [20, 177] greater in VPM neurons (202 ms, *SD* 119) than VPL neurons (104 ms, *SD* 92, *p* = 0.016; **Figure 4E**). Each of these effects indicate decreased spike firing in VPM neurons in response to depolarizing stimuli suggesting that VPM neurons may have reduced excitability relative to VPL neurons.

**Figure 4.**
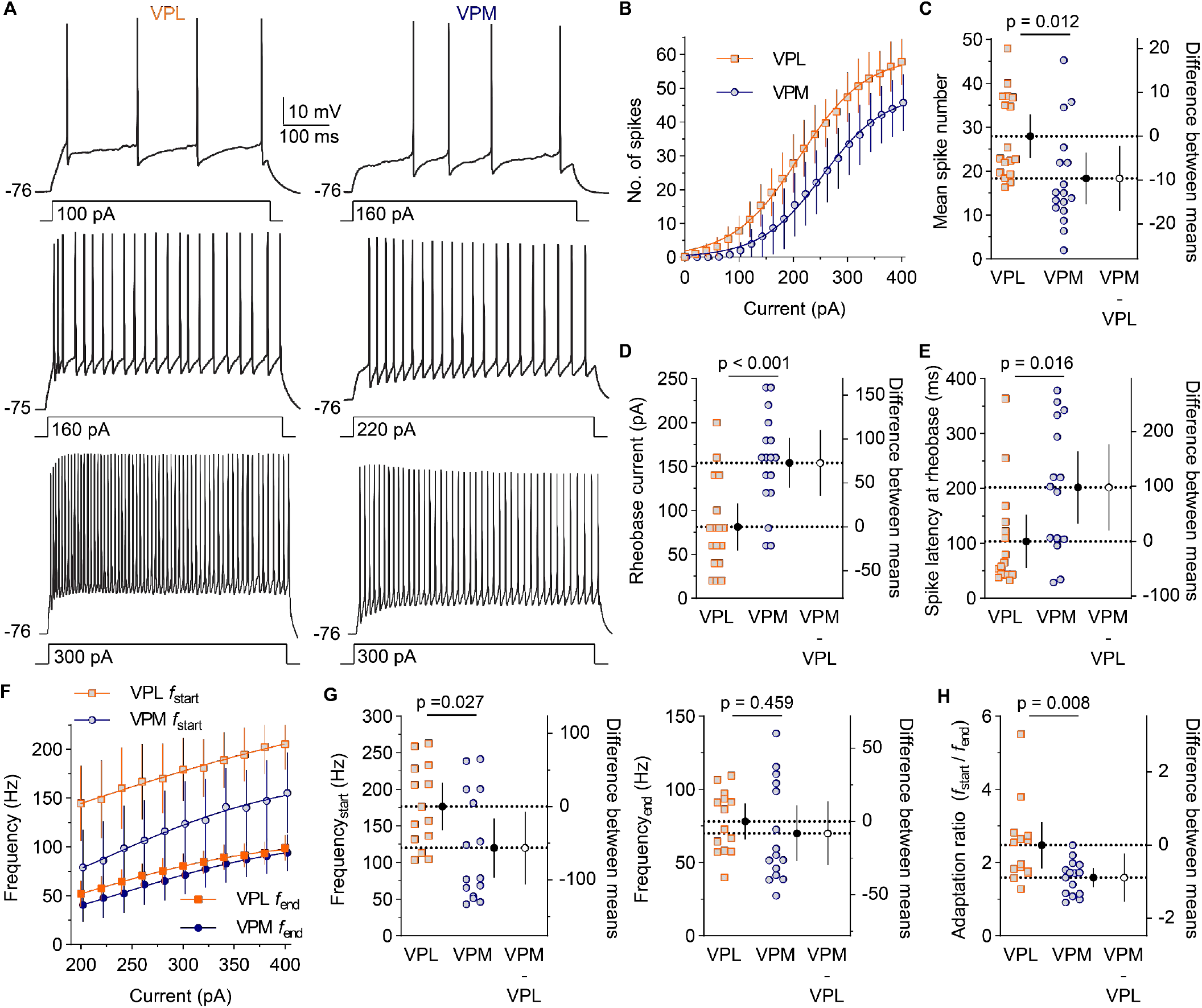
VPL neurons have enhanced spiking and spike adaptation during depolarization. **A.** Spikes were elicited by 500 ms depolarizing current injections (0 – 400 pA). **B – H**. Point estimates and error bars are group means with 95% CI. **B**. Spike number was plotted vs. current amplitude. VPL: *n* = 17 cells, 11 mice. VPM: *n* =17 cells, 12 mice. Mixed-effects model: main effect *F* (1, 32) = 7.054, *p* = 0.012; interaction *F* (20, 640) = 2.884, *p* < 0.001. (**C**) The overall mean spike number across current injections (main effect from panel B), (**D**) rheobase current (Welch’s t test, *t* = 3.962, *df* = 31.90), and (**E**) spike latency at rheobase (Welch’s t test, *t* = 4.189, *df* = 21.25) were plotted as individual data points for each cell alongside the group mean (black circle, dashed line). The difference between means (VPM – VPL) was plotted (open circle) on the right y-axis. **F**. *f*_start_ and *f*_end_ were plotted versus current amplitude. VPL: *n* = 14 cells, 9 mice. VPM: *n* =15 cells, 10 mice. Mixed-effects model: *f*_start_, main effect *F* (1, 27) = 5.376, *p* = 0.027, interaction *F* (10, 270) = 0.829, *p* = 0.601; *f*_end_, main effect *F* (1, 27) = 0.564, *p* = 0.459, interaction *F* (10, 270) = 2.041, *p* = 0.030. (**G**) The mean *f*_start_ and *f*_end_ (main effects from panel F) as well as (**H**) adaptation ratio (mixed-effects model: main effect, *F* (1, 27) = 8.226; interaction, *F* (10, 270) = 2.939, *p* = 0.002) across all current injections were plotted as in panels C – E.

Spike adaptation is a reduction in AP frequency over a sustained period of depolarization. To test for differences in spike adaptation, we quantified the frequency of the initial three spikes (*f*_start_) and the last two spikes (*f*_end_) across current injections between 200 – 400 pA, and then computed an adaptation ratio (*f*_start_ / *f*_end_). At the start of the spike train, the overall mean frequency across current injections was 57 Hz [-107, -7] lower in VPM neurons (120 Hz, *SD* 73) relative to VPL neurons (177 Hz, *SD* 56, *p* = 0.027); whereas, the mean frequency at the end of the spike train was 8 Hz [-30, 14] lower in VPM neurons (70 Hz, *SD* 34) than VPL neurons (78 Hz, *SD* 21, *p* = 0.459; **Figure 4F,G**). The mean adaptation ratio was reduced by 0.9 [-1.5, -0.3] in VPM neurons (1.6, *SD* 0.5) compared to VPL neurons (2.5, *SD* 1.1; *p* = 0.008; **Figure 4H**). Together, these data are consistent with reduced spiking upon depolarization in VPM neurons, but enhanced spike adaptation in VPL neurons during sustained depolarization.

### Hyperpolarization-induced rebound bursting and sag potential differ between VPM and VPL neurons

Hyperpolarization-induced rebound bursting in both VPL and VPM neurons contributes to thalamic oscillations and impacts sensory signal propagation to the cortex. Recent evidence revealed distinct, nucleus-specific nRT connectivity with VPL and VPM neurons indicating the two thalamic regions may contribute distinctly to circuit-wide oscillations (Clemente-Perez et al., 2017). Here, we directly compared properties of hyperpolarization-induced rebound bursting in VPL and VPM neurons (**Figure 5A**). Across all current injections, the overall mean spike number per burst was 5 spikes (*SD* 2) for VPM neurons and 6 spikes (*SD* 3) for VPL neurons, which is a mean difference of -1 spike [-3, 1], *p* = 0.307 (**Figure 5B,C)**. The mean spike frequency within bursts was 27 Hz [-16, 70] higher in VPM neurons (268 Hz, *SD* 32) than VPL neurons (241, *SD* 63, *p* = 0.094; **Figure 5D,E**). The mean spike adaptation within bursts was 1.9 (*SD* 0.2) in VPM neurons and 1.7 (*SD* 0.2) in VPL neurons, which is a mean difference of 0.2 [0.0, 0.4], *p* = 0.061 (**Figure 5F**). These data suggest that VPL and VPM neurons fire rebound bursts with similar spike number and frequency. Next, we compared the shape of APs within bursts from VPL and VPM neurons. The overall mean amplitude of burst APs elicited by the maximum hyperpolarizing current injection (400 pA) was 6.3 mV [-11.7, -1.1] lower in VPM neurons (61.3 mV, *SD* 7) than VPL neurons (67.7 mV, *SD* 5; *p* = 0.021; **Figure 5G**). The overall mean AP half width was 0.1 ms [-0.3, 0.4] longer in VPM neurons (1.5 ms, *SD* 0.4) than VPL neurons (1.4 ms, *SD* 0.4; *p* = 0.681; **Figure 5H**). These data are consistent with no substantial changes in AP shape and frequency within rebound bursts as confidence intervals for all measures include or approach zero, which raises uncertainty about the potential biological significance of the minimal effects that showed statistical significance such as AP amplitude.

**Figure 5.**
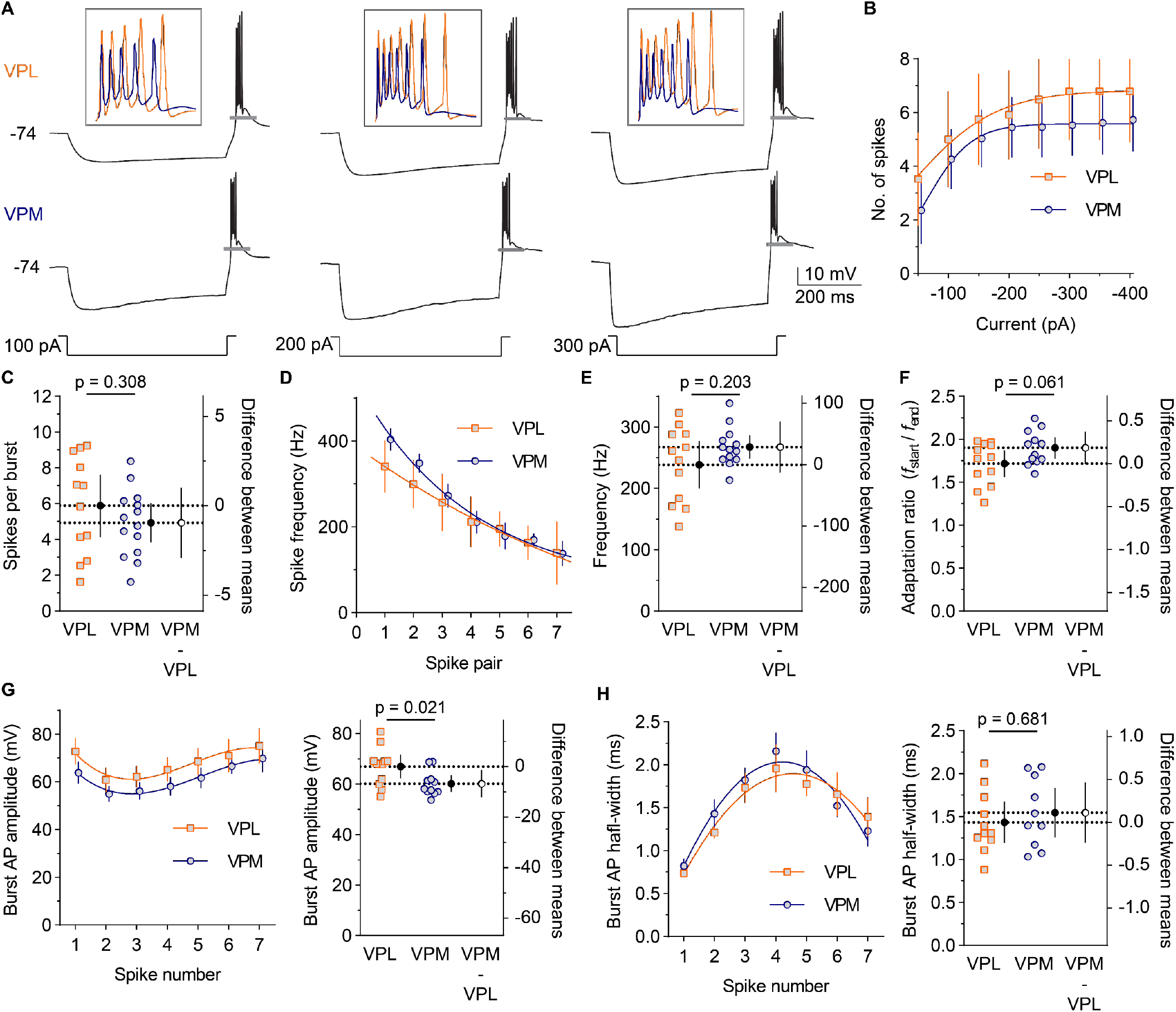
VPL and VPM neurons have similar rebound burst spike number and frequency. **A.** Bursts were elicited by 500 ms hyperpolarizing current injections. **B – H**. Point estimates and error bars are group means with 95% CI. **B**. Spike number per burst was plotted versus current amplitude. VPL: *n* = 12 cells, 8 mice. VPM: *n* =14 cells, 8 mice. Mixed-effects model: main effect *F* (1, 24) = 1.088, interaction *F* (7, 168) = 1.035, *p* = 0.409. **C**. The overall mean spike number (main effect from panel B) was plotted as individual data points for each cell alongside the group mean (black circle, dashed line) on the left y-axis. The difference between means (VPM – VPL) was plotted on the right y-axis (open circle). **D**. The spike frequency was plotted for each spike pair within bursts. Mixed-effects model: main effect *F* (1, 24) = 1.718, interaction *F* (7, 99) = 3.635, *p* = 0.002. The overall mean (**E**) frequency (main effect from panel D) and (**F**) adaptation ratio (main effect *F* (1, 22) = 3.896, *p* = 0.061, interaction *F* (5, 77) = 2.703, *p* = 0.027) were plotted as in panel C. **G**. AP amplitude (main effect *F* (1, 21) = 6.211, interaction *F* (6, 101) = 0.759, *p* = 0.604) and (**H**) half width (main effect *F* (1, 21) = 0.175, interaction *F* (6, 101) = 0.531, *p* = 0.783) were plotted for each spike (left). VPL: *n* = 12 cells, 8 mice. VPM: *n* = 11 cells, 7 mice. AP amplitude and half-width (main effects) were averaged across spikes and plotted as described for panel C.

In addition, we compared the burst latency, rebound depolarization duration, and sag potential of the voltage responses to hyperpolarization in VPL and VPM neurons. To do so, we analyzed voltage responses to a single current injection (300 pA), which allowed us to measure how these parameters were correlated within individual responses. The mean latency between removal of the hyperpolarizing current and rebound burst firing was 3.7 ms [-5.8, -1.5] shorter in VPM neurons (11.5 ms, *SD* 1.7) compared to VPL neurons (15.1 ms, *SD* 3.0, *p* = 0.002; **Figure 6A,B)**. The rebound depolarization period was 0.4 s [0.3, 0.7] longer in VPM neurons (1.4 s, *SD* 0.2) compared to VPL neurons (1.0 s, *SD* 0.2, *p* < 0.001; **Figure 6C,D,F**). Interestingly, the sag potential amplitude during hyperpolarization was 3.8 mV [1.6, 5.8] larger in VPM neurons (12.2 mV, *SD* 2.6) compared to VPL neurons (8.6 mV, *SD* 2.3, *p* = 0.010; **Figure 6E,G**). Increased sag potential could indicate that VPM neurons have increased hyperpolarization-activated depolarizing currents (I_h_), which is consistent with a more robust rebound depolarization and reduced burst latency. Indeed, sag potential amplitude was negatively correlated with burst latency for VPL neurons (*r* = -0.760, 95% CI [-0.929, -0.329], *R^2^* = 0.577, *p* = 0.004) and VPM neurons (*r* = -0.827, 95% CI [-0.954, -0.452], *R*^2^ = 0.684, *p* = 0.002; **Figure 6H**). In addition, sag potential was positively correlated with rebound depolarization duration for VPL neurons (*r* = 0.782, 95% CI [0.377, 0.936], *R*^2^ = 0.611, *p* = 0.003) and VPM neurons (*r* = 0.829, 95% CI [0.457, 0.954], *R*^2^ = 0.688, *p* = 0.002; **Figure 6I**). Together, these results suggest VPL and VPM neurons have minimal differences in spike frequency or shape within rebound bursts, but may respond differently to hyperpolarizing stimuli, which results in faster burst generation in VPM neurons upon recovery from hyperpolarization.

**Figure 6.**
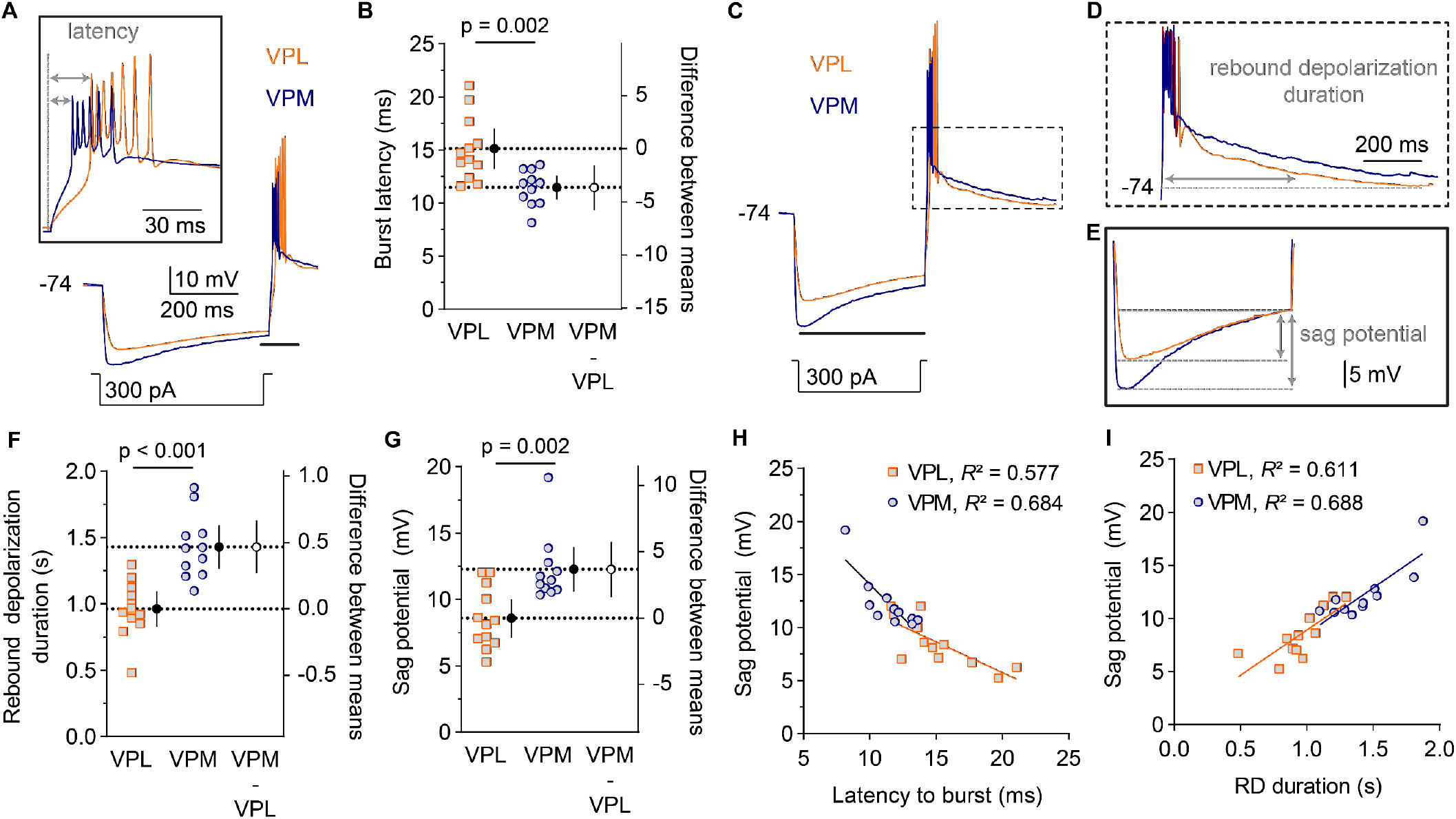
VPM neurons have reduced burst latency relative to VPL neurons. **A.** The representative traces show voltage responses to 300 pA hyperpolarizing current injections. Inset: the rebound burst region above the black line was expanded to visualize the period between removal of hyperpolarization and the end of rebound burst firing. **B**. The latency from removal of the hyperpolarizing current to the first spike was plotted as individual data points for each cell alongside the group mean (black circle, dashed line) with 95% CI on the left y-axis. The difference between means (VPM – VPL) was plotted (open circle) with the 95% CI on the right y-axis. VPL: *n* = 12 cells, 7 mice. VPM: *n* = 12 cells, 7 mice. Welch’s t test: *t* = 3.651, *df* = 17.48. **C.** A representative trace was expanded to visualize (**D**) the full rebound depolarization period (boxed region in panel C) and (**E**) the sag potential during hyperpolarization (region above black line in panel C). Welch’s t tests were used to compare the (**F)** rebound depolarization duration (*t* = 4.907, *df* = 19.91. and (**G**) sag potential (*t* = 3.645, *df* = 20.26), which were plotted as described for panel B. P values were reported above each plot. The sag potential amplitude for each cell was plotted vs. (**H**) burst latency and (**I**) rebound depolarization (RD) duration. Data were fit by linear regression and analyzed with Pearson’s correlation coefficients.

## Discussion

VPL and VPM thalamus comprise two neighboring thalamocortical neuron populations that are part of parallel pathways carrying somatosensory information from the body and head, respectively. Here, we revealed several physiological properties that differ between VPL and VPM neurons indicating that important functional differences may regulate how these distinct ascending sensory signals are transmitted or processed in the thalamus. Specifically, analysis of spontaneous synaptic transmission suggests substantial differences in both excitatory and inhibitory synaptic input to VPL and VPM neurons. Tonic and burst firing properties also differed between VPL and VPM neurons, which may contribute to their distinct physiological roles in corticothalamic network function. Furthermore, this work indicates that neurons in the ventrobasal thalamus, also called the ventral posterior thalamus, are more heterogeneous than previously recognized, thus highlighting the importance of applying cell-type- or nucleus-specific methodologies when investigating thalamic circuitry.

In this study, we observed that VPM neurons had reduced frequency and amplitude of glutamatergic synaptic input compared to VPL neurons suggesting that excitatory drive to VPM neurons may be weaker than VPL neurons. Somatosensory information is carried via glutamatergic projections through the medial lemniscus and spinothalamic tracts to VPL neurons and the trigeminothalamic tract to VPM neurons. VPL and VPM neurons also receive descending glutamatergic input from layer 6 corticothalamic neurons. However, it remains unknown whether differences in the innervation and/or function of cortical or sensory inputs underlie the reduced excitatory synaptic input observed in VPM neurons. To our knowledge, the density and convergence of ascending and descending projections have not been directly compared between VPL and VPM neurons, but it is possible that individual VPM neurons receive fewer inputs due to less convergence of sensory or cortical afferents compared to the VPL. Alternatively, it is possible that there are fewer neurons projecting to the VPM thalamus compared to the VPL or that the neurons projecting to VPM thalamus have lower spontaneous activity resulting in decreased frequency of glutamatergic synaptic events. Synapse strength may differ between VPL and VPM neurons due to different presynaptic release properties at their anatomically distinct inputs or distinct postsynaptic receptor expression. A limitation of this study is that we have not used synaptic stimulation via ascending or descending inputs to evoke VPL and VPM neuron activity, which would be necessary to determine how these differences in synapse physiology might affect excitatory drive to the thalamus.

GABAergic neurons in the nRT provide the major inhibitory input to VPL and VPM neurons. Recent work suggests that PV-positive nRT neurons project to both VPM and VPL, whereas SOM-positive nRT neurons project only to VPL neurons (Clemente-Perez et al., 2017). Furthermore, the nRT-VPM functional connectivity may be stronger than nRT-VPL connectivity. Our investigation of spontaneous inhibitory synaptic transmission indicated minimal differences in overall inhibitory synaptic event frequency; however, our data were consistent with substantially larger amplitude bursts of inhibitory synaptic input in VPM neurons relative to VPL neurons. When nRT neurons are hyperpolarized they exhibit high-frequency burst firing in response to excitatory input, and PV-positive neurons have stronger burst firing properties than SOM-positive nRT neurons including more spikes per burst (Clemente-Perez et al., 2017). Therefore, the enhanced bursts of inhibitory input observed in VPM neurons could be due to stronger input from PV-positive neurons compared to VPL neurons, which would be consistent with previous work indicating that the nRT-VPM loop is more rhythmogenic (Clemente-Perez et al., 2017). However, the impact of these collective differences in inhibitory and excitatory synapse function on CT-nRT-VPL/VPM feed-forward inhibition and the propagation of sensory information through the VPL and VPM thalamus remains unclear.

In terms of cellular excitability, our study suggests that depolarization-induced firing of VPL neurons was enhanced compared to VPM neurons, including increased number of spikes, reduced spike latency, and reduced rheobase. The increased firing upon depolarization and reduced latency to the first spike in VPL neurons compared to VPM neurons suggests that VPL neurons may have increased excitability in response to depolarizing excitatory synaptic input. The mechanism mediating this difference in depolarization induced spiking remains unclear as there were no significant differences in membrane properties between the two cell populations. The rising phase of the AP in VPL and VPM neurons appeared similar, but voltage-gated channel expression and function have not been compared between VPL and VPM neurons; therefore, we cannot rule out differences in expression or subcellular localization of voltage-gated sodium or calcium channel isoforms. Indeed, haploinsufficiency of Na_V_1.1 in a Dravet syndrome mouse model led to opposing changes in VPL and VPM cellular excitability suggesting that voltage-gated sodium channels may have differential expression or function in the two populations (Studtmann et al., 2022). Understanding the precise mechanisms underlying differences in VPL and VPM depolarization-induced firing will require a systematic investigation of the membrane channels contributing to cellular excitability in each neuron population.

VPL neurons also exhibited enhanced spike frequency adaptation compared to VPM neurons, which may suggest that a unique activity-dependent mechanism is contributing to VPL firing across sustained depolarization. Spike frequency adaptation in the VB thalamus may be mediated by anoctamin-2 (ANO2), a calcium-activated chloride channel (Ha & Cheong, 2017; Ha et al., 2016). ANO2 becomes activated by calcium influx through voltage-gated calcium channels during sustained depolarization, and the resulting influx of chloride ions hyperpolarizes the neuron, thereby reducing the propensity to spike (Ha & Cheong, 2017). However, the expression and function of ANO2, or other channels implicated in spike frequency adaptation such as calcium-activated potassium channels (e.g. SK channels), have not been investigated in VPL and VPM neurons independently. ANO2 and SK channels also contribute to the AHP period following spike firing in VB neurons (Ha & Cheong, 2017; Ha et al., 2016). Interestingly, we observed a larger AHP in VPL neurons relative to VPM neurons, which further suggests there may be differences in these calcium-activated currents in the two neuron populations. Physiologically, spike frequency adaptation in VB neurons has been proposed as a self-inhibiting mechanism, limiting the propagation of information during excessive activation (Cheong et al., 2011; Ha et al., 2016; Landisman & Connors, 2007). Therefore, differences in spike frequency adaptation between VPL and VPM neurons could suggest that the thalamus may differentially regulate propagation of somatosensory information from the body and head.

Furthermore, we observed interesting differences in burst firing properties of VPL and VPM neurons following removal of hyperpolarizing currents. Feedforward CT-nRT-VPL/VPM inhibition provides a robust hyperpolarizing stimulus to VPL and VPM neurons, which then respond to subsequent depolarization with a low-threshold calcium spike, mediated by T-type calcium channels, and high-frequency rebound burst firing. Recurrent rebound bursts between the nRT and the VB thalamus generate oscillations, which underlie important physiological processes such as sleep spindles. It has been postulated that the nRT-VPM loop may be more important in intra-thalamic oscillation generation and maintenance than the nRT-VPL loop, and, as stated earlier, this may be due to the stronger coupling between PV-positive nRT neurons and VPM neurons (Clemente Perez et al., 2017). However, VPM neurons may have intrinsic burst firing properties that contribute to their role in thalamic oscillations. We observed no substantial differences in spike number or frequency within bursts, but VPM neurons did exhibit a substantially shorter latency to burst firing upon removal of hyperpolarization and a greater sag potential during hyperpolarization. While burst firing is mediated, in large part, by T-type calcium channels, HCN channels mediate a hyperpolarization-activated cation current (I_h_) that also regulates burst firing properties in TC neurons (Llinas & Steriade, 2006). I_h_ is responsible for the sag potential during sustained hyperpolarization, and sag potential amplitude has been shown to negatively correlate with burst latency, but have no correlation to the number of spikes per burst (Desai & Varela, 2021). Thus, our observation of greater sag potentials in VPM neurons could be caused by increased I_h_ relative to VPL neurons, which is consistent with reduced latency to burst firing in VPM neurons. nRT and VB neurons express different T-type calcium channel isoforms with nRT neurons expressing CaV3.2 and CaV3.3, while VB neurons express predominantly CaV3.1 (Astori et al., 2011; Talley et al., 1999). In addition to T-type calcium channels, both VB and nRT neurons express HCN channels that are activated at hyperpolarized potentials and further contribute to rebound bursting (Abbas et al., 2006; Zobeiri et al., 2019). Yet, CaV and HCN channel function has not been investigated in the VPL and VPM individually. Elucidating the specific mechanisms creating distinct tonic and burst firing properties will require a direct comparison of ion channel expression and function in the two cell populations.

From a physiological perspective, the differences in synaptic function, depolarization-induced firing, and rebound bursting indicate that the VPL and VPM may process somatosensory information differently. There is likely an evolutionary advantage to processing somatosensory information from the body and head through parallel circuits with distinct physiology, though at this time we can only speculate what that advantage might be. Both the number and frequency of spike trains encode critical information that is propagated from the VPL and VPM to the somatosensory cortex (Kenshalo et al., 1980; Martin et al., 1996). Specifically, VPL neurons have been shown to increase their frequency of output in response to increasingly noxious stimuli, but not innocuous sensory input (Kenshalo et al., 1980; Martin et al., 1996). Thus, it is possible that VPL neurons have a different sensitivity to incoming signals from noxious stimuli compared to VPM neurons. The enhanced spike frequency adaptation in the VPL may indicate they are more likely to slow their response to repeated incoming sensory signals than their VPM counterparts. A limitation of this study is that it is difficult to interpret how these changes in excitability impact overall sensory processing as this would require evaluating VPL and VPM responses to ascending input. It is possible that neurons within the ascending somatosensory tracts to the VPL and VPM have unique cellular properties and excitability. Therefore, the VPL and VPM could receive ascending input with distinct patterns or strengths that also shape how they process and relay sensory signals. Thus, it is crucial to understand the integration of ascending input as well as descending cortical input to both nuclei in order to interpret how their unique excitability contributes to overall sensory processing.

The data presented here, as well as recent evidence regarding distinct inputs to the VPL and VPM, strongly suggest that the VPL and VPM form functionally distinct microcircuits within the broader somatosensory CT circuit (Clemente Perez et al., 2017). Understanding the functional differences between these two parallel pathways is not only important for better understanding of a healthy circuit, but may also have critical implications for a variety of diseases including epilepsy, chronic pain, and sleep disorders. Investigations into pathological function within the somatosensory CT circuit have historically treated the VB thalamus as a uniform complex, potentially obscuring nucleus-specific effects in the VPL and VPM. This is therapeutically important as identifying the specific dysfunction within each nucleus may reveal previously unidentified disease mechanisms and therapeutic opportunities. Further investigation is required to uncover the molecular mechanisms through which these two microcircuits process sensory signals and generate circuit-wide rhythmogenesis as well as how they may be uniquely suited for therapeutic targeting.

## Supplemental Figures

**Figure S1.**
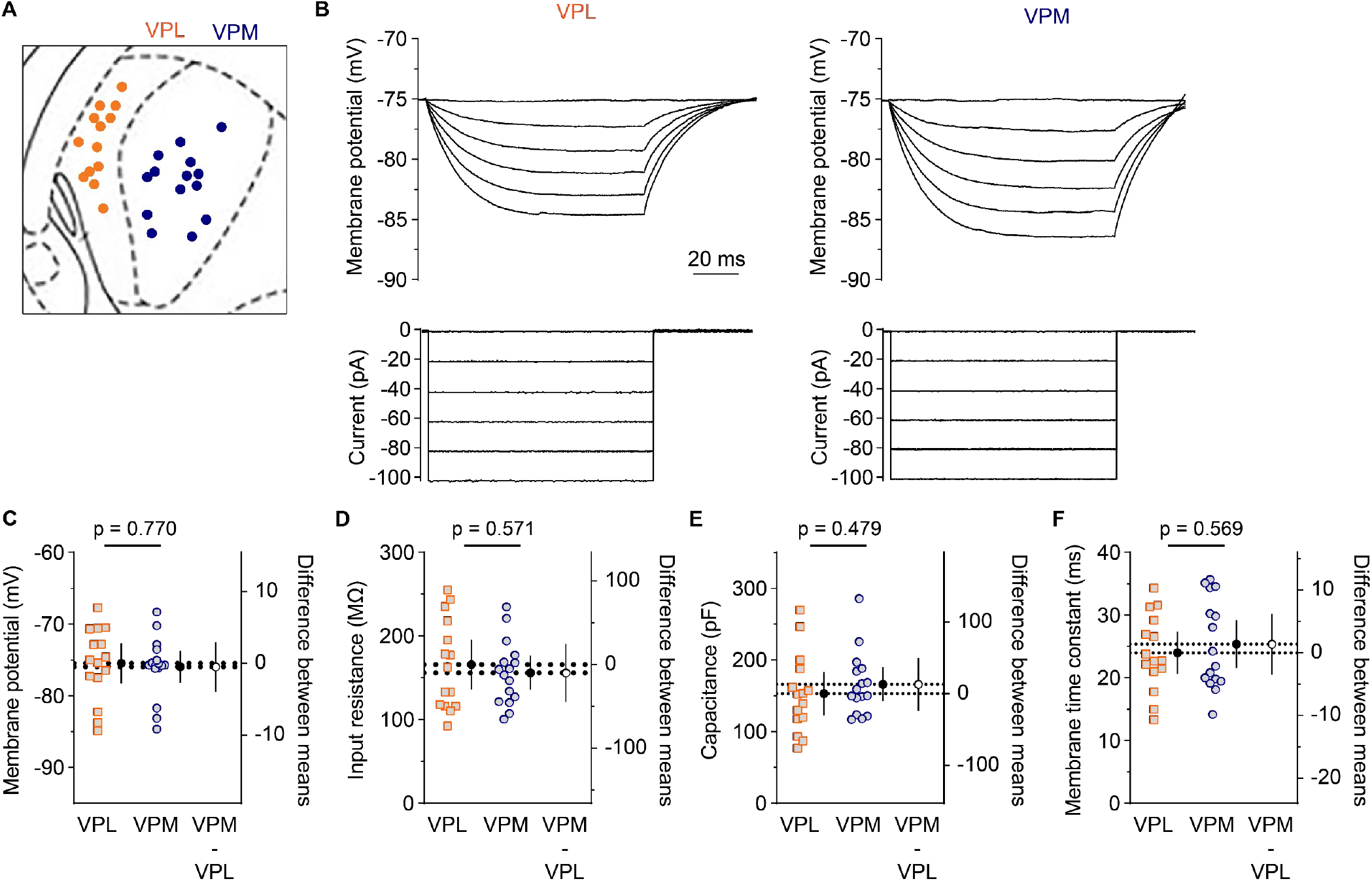
Intrinsic membrane properties of VPL and VPM neurons. **A.** The location of neurons used for all current clamp recordings (Figures 3 – 6) were mapped onto an image of the VPL and VPM thalamus. **B.** Intrinsic membrane properties were analyzed using the voltage responses to 100 ms hyperpolarizing current injections (0 – 100 pA). **C.** Resting membrane potential, (**D**) input resistance, (**E**) cell capacitance, and (**F**) membrane time constant were quantified and plotted as individual data points for each cell alongside group means with 95% CI on the left y-axis. The difference between the means (VPM – VPL) were plotted with the 95% CI on the right y-axis. Groups were compared by Welch’s t tests, and p values were reported above each plot.

